# BROAD-NESS uncovers dual-stream mechanisms underlying predictive coding in auditory memory networks

**DOI:** 10.1101/2024.10.31.621257

**Authors:** Leonardo Bonetti, Gemma Fernández-Rubio, Mathias H. Andersen, Chiara Malvaso, Francesco Carlomagno, Claudia Testa, Peter Vuust, Morten L. Kringelbach, Mattia Rosso

**Author notes:** Corresponding authors (L.B.); (M.R.).

## Abstract

Auditory memory enables the recognition of sound sequences by integrating sensory input with memory traces and predictive mechanisms. While predictive coding has been proposed as a key framework for this process, the large-scale brain networks supporting it remain poorly understood. Using BROADband brain Network Estimation via Source Separation (BROAD-NESS), we provide three fundamental insights into the neural organisation of auditory memory and predictive coding. First, auditory cortices participate in two distinct, orthogonal whole-brain networks, expanding on the conventional view focused on forward and backward information flow between single brain regions. One network involves the medial cingulate gyrus, while the other integrates prefrontal and hippocampal regions, inferior temporal cortex, and insula. Second, we present evidence for a dual-stream mechanism in auditory memory recognition, which both parallels and diverges from the well-established dual-stream hypothesis in vision. Third, predictive coding in conscious auditory memory is supported by large-scale networks generating confirmed predictions and prediction errors. While previous studies examined predictive coding in isolated regions or pairwise connections, our findings reveal how whole-brain networks coordinate these processes, highlighting fine-grained spatial gradients and distinct temporal dynamics. These findings enhance our understanding of auditory perception, memory, and prediction, as well as their underlying basis in whole-brain dynamical networks.

## Introduction

Predictive coding is a fundamental principle of brain function, proposing that perception and cognition rely on hierarchical predictions to minimise discrepancies between expected and incoming sensory information^1,2,3^. This framework has been extensively investigated in relation to automatic sensory processes, often indexed by event-related potential/field (ERP/F) components such as N100, mismatch negativity (MMN), and error-related negativity (ERAN)^4–12^. These studies have demonstrated that such components are automatically elicited in response to deviations in visual and auditory stimuli, including changes in expected image and sound features, variations in the probability of sound occurrence (N100, MMN)^4,5,6,7,8,9,10^, and musical harmonic properties (ERAN)^11,12^. In addition, considerable research has explored conscious predictive coding, where individuals explicitly anticipate upcoming stimuli. Most studies in this domain have examined its neural correlates within decision-making^13^, memory-based predictions^14,15^, or the interplay between conscious and automatic predictive processes^16^.

As in many areas of cognitive neuroscience, predictive coding research has also progressively transitioned from activation-based approaches towards connectivity analyses. This shift was driven by mounting evidence that the brain operates as a complex system, where regions interact within distributed networks rather than functioning in isolation^17,18^. Previous research has used various metrics to assess functional connectivity, with dynamic causal modelling (DCM) emerging as the primary method for inferring connectivity between a limited set of brain regions in the context of predictive coding^19^. For instance, DCM has been widely used to investigate the hierarchical organisation of predictive coding during automatic processing, revealing the flow of information from the primary auditory cortex to the superior temporal gyrus and inferior frontal gyrus in the MMN context^20,21^. Similarly, DCM studies in vision have shown that imprecise target motion increases the gain of superficial pyramidal cells in visual cortex^22,23^. Beyond automatic processing, DCM has also been applied to social communication, demonstrating that social demands preferentially modulate backward connections from the medial prefrontal cortex (MPFC) rather than forward connections from the superior occipital gyrus (SOG) and medial temporal gyrus (MTG) to the MPFC^24^. Additionally, studies combining DCM with transfer entropy have provided further insights into predictive coding in aging, multisensory integration, and conscious processing, showing that older adults rely more on cross-modal predictive neural templates^25^.

Despite the substantial focus on predictive coding, long-term memory and predictive processes for temporally unfolding information have historically received little attention. This was rather surprising, given that most stimuli humans encounter are inherently structured over time. Thus, as this realisation emerged, the topic has, in recent years, attracted significant interest. For example, Albouy and colleagues^26^ investigated the neural mechanisms underlying memory retention for temporal sequences, demonstrating that theta oscillations in the dorsal stream predict auditory working memory performance. Similarly, Quiroga-Martinez and colleagues^27^ showed that predictive processes and the manipulation of temporally structured sound information can be directly decoded from non-invasive neurophysiological data, revealing a widespread network encompassing the auditory cortices and medial temporal lobe.

Along this line, we conducted a series of studies using diverse tools to estimate dynamic functional connectivity during the long-term encoding and recognition of temporal sequences. This revealed a large network of brain regions involved in processing both memorised and novel musical sounds, extending from the auditory cortex to the medial cingulate, inferior temporal cortex, insula, frontal operculum, and hippocampus, and showing precise dynamical changes in response to each sound of the sequences^28–30^. Moreover, we discovered that the activity of this brain network was modulated by musical complexity^31^ and individual cognitive differences^32^. In a subsequent study, following common practice in predictive coding research, we also employed DCM to examine directional functional connectivity between six predefined large-scale regions of interest (ROIs) during long-term recognition and prediction of musical sequences. This analysis provided the strongest model evidence for a network in which information flowed from auditory cortices to higher-order regions such as the hippocampus and ventromedial prefrontal cortex (vmPFC) in both forward and backward directions^33^. However, despite being informative, DCM typically involves comparing a hypothesised model against only a limited set of alternatives. Thus, to complement this approach, we recently extended our investigation using a novel multivariate framework, Directed Multiplex Visibility Graph Irreversibility (DiMViGI), which systematically assessed all possible ROI configurations to identify those exchanging the highest amount of information. This revealed distinct peak interactions between the auditory cortex, hippocampus, and vmPFC within each hemisphere, indicating a functionally segregated yet interactive network supporting long-term recognition of temporal information^34^.

Although these studies significantly advanced our understanding of auditory predictive coding, they remained constrained to a predefined set of ROIs, fundamentally limiting a comprehensive characterisation of whole-brain connectivity and raising a fundamental question: How are auditory cortices embedded within large-scale brain networks, potentially engaging in multiple concurrent computations to support predictive and memory processes?

To address this, we draw on previous studies employing linear decomposition techniques to disentangle overlapping brain processes. These methods have been successfully applied in fMRI research, particularly through Principal Component Analysis (PCA) and Independent Component Analysis (ICA)^35–39^ (for a review, see^40^). However, their application to predictive and memory processing has been limited, likely due to fMRI’s low temporal resolution^41^, which does not allow the computation of reliable covariance matrices for rapid events. By contrast, neurophysiological recordings (EEG/MEG) offer superior temporal resolution, enabling the investigation of rapid covariance patterns over brief time windows, as required for predictive coding and memory studies. Nonetheless, traditional EEG/MEG analyses have primarily applied PCA at the scalp level^42–46^, limiting insights into source-level networks (for a review, see^47^). One exception is a study that used ICA to derive resting-state brain networks from MEG data^48^, revealing configurations similar to those observed in fMRI. However, this study focused solely on canonical frequency bands in the resting state, offering little insight into event-related network dynamics and their variance contributions.

To overcome these limitations, we recently developed Network Estimation via Source Separation (NESS), a multivariate framework that applies linear decomposition algorithms to estimate brain networks from source-reconstructed MEG data^49^. Unlike previous methods, NESS directly identifies simultaneous brain networks, providing their explained variance, fine-grained spatial configurations for physiological interpretation, and activation time series for further analysis. Its frequency-resolved variant, FREQ-NESS^50^, was recently shown to successfully distinguish concurrent, frequency-specific networks during resting state as well as their reorganisation in response to auditory stimuli.

Here, we extend this approach by introducing BROAD-NESS, a variant of the NESS framework that applies PCA to broadband MEG source-reconstructed voxel time series recorded during event-related tasks. Using BROAD-NESS on MEG data from a musical sequence recognition task, we aimed to determine how auditory cortices are embedded in large-scale whole-brain networks and whether they engage in multiple concurrent computations supporting predictive coding and memory processes. This work represents a crucial step towards bridging the gap in our understanding of conscious predictive processes at a whole-brain network level.

## Results

### Overview of the experimental design and MEG source reconstruction

Eighty-three participants engaged in an old/new auditory recognition task during magnetoencephalography (MEG) recordings (**Figure 1a-b**). Following the learning of a brief musical piece (see **Figure S1**), participants were presented with 135 five-tone musical sequences, each lasting 1750 ms. They were asked to indicate whether each sequence belonged to the original music (‘memorised’ sequence, denoted as M, old) or represented a varied sequence (‘novel’ sequence, denoted as N, new) (Figure 1a). Among the presented sequences, 27 were directly extracted from the original musical piece, while 108 were variations. These variations were categorised based on the number of musical tones altered following the first tone (NT1), second tone (NT2), third tone (NT3), or fourth tone (NT4) (see **Figure S2**).

**Figure 1:**
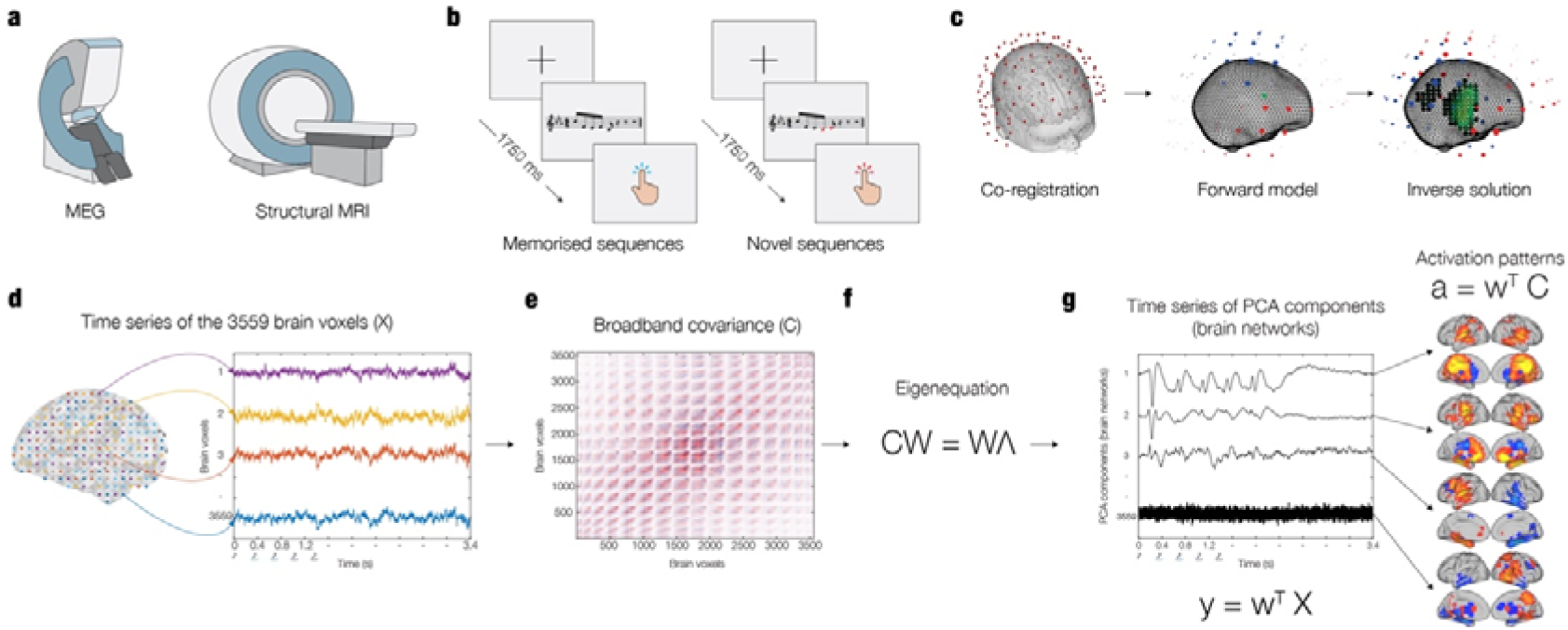
BROADband network estimation via source separation (BROAD-NESS) The figure provides an overview of the BROAD-NESS methodology. **a -** MEG was used to collect neurophysiological data while participants performed an auditory old/new recognition task. **b -** Five-tone auditory sequences, presented in random order, were classified by participants as either "old" (memorised sequences, M) or "new" (novel sequences, N) using button presses. **c -** Co-registration was performed between MEG data and individual MRI anatomical scans. To reconstruct the neural sources which generated the signal recorded by the MEG, a single shell forward model was used. The inverse solution was estimated through beamforming. **d -** The source reconstruction yielded time series data for 3,559 brain voxels based on an 8-mm grid brain parcellation. **e -** The covariance matrix (C) was computed from the broadband voxel data matrix. **f** – Principal Component Analysis (PCA) was computed by solving the eigenequation CW = WΛ for the eigenvectors (W) to find the weighted combinations of brain voxels to orthogonal components, which reflected brain networks; the associated eigenvalues (Λ) express the amount of variance explained by each network component. **g -** Network activation time series (y) were derived by applying the spatial filters (W) to the voxel data matrix (X). The same filters were applied to the covariance matrix (C) to compute the spatial activation patterns (a), representing the components’ projections in voxel space.

After performing the pre-processing of the MEG data (see Methods for details), we computed source reconstruction. We employed a single-shell forward model alongside a beamforming approach as the inverse solution, using an 8-mm grid that corresponded to 3,559 brain voxels (**Figure 1c-d**). This procedure produced a time series for each reconstructed brain voxel returning a matrix of 3,559 brain voxels x time-points, independently for each participant and experimental condition.

### PCA, Monte Carlo Simulations (MCS) and statistics

PCA was conducted on the reconstructed time series of the 3,559 brain voxels, averaged across participants and conditions, to identify simultaneously operating broadband brain networks (**Figure 1e-f-g**). To assess significant Principal Components (PCs), a Monte Carlo Simulation (MCS) approach was used by performing PCA on data randomised across the temporal dimension for 100 permutations. Significant principal components from the original data were defined as those explaining more variance than the maximum variance accounted for by the first component across the 100 permutations of the randomised data (see Methods for details). Remarkably, the variance explained by the principal components in the randomised data remained rather consistent across all permutations, suggesting that a conventional set of 100 permutations may not be necessary since a reduced number of permutations (e.g., 50 or even as few as 10) would indeed yield equivalent results.

This procedure identified six significant PCs, which accounted for the following percentages of variance: 71.75%, 16.32%, 4.16%, 1.30%, 0.99%, 0.92%. Each component was interpreted as a distinct brain network based on its spatial activation pattern, which was obtained by multiplying the component’s eigenvector by the covariance matrix of the data. Unlike FREQ- NESS, which was based on GED, the weights and spatial activation patterns produced by PCA were equivalent in demonstrating the relative contribution of each brain voxel to the network.

As illustrated in **Figure 2**, the first identified brain network comprised the auditory cortices and the medial cingulate gyrus, while the second one included the auditory cortices, anterior cingulate gyrus, ventromedial prefrontal cortex, hippocampal regions, and insula. The remaining four networks were characterised by similar spatial maps; however, they were deemed less relevant as they explained considerably less variance (**Figure S3**).

**Figure 2:**
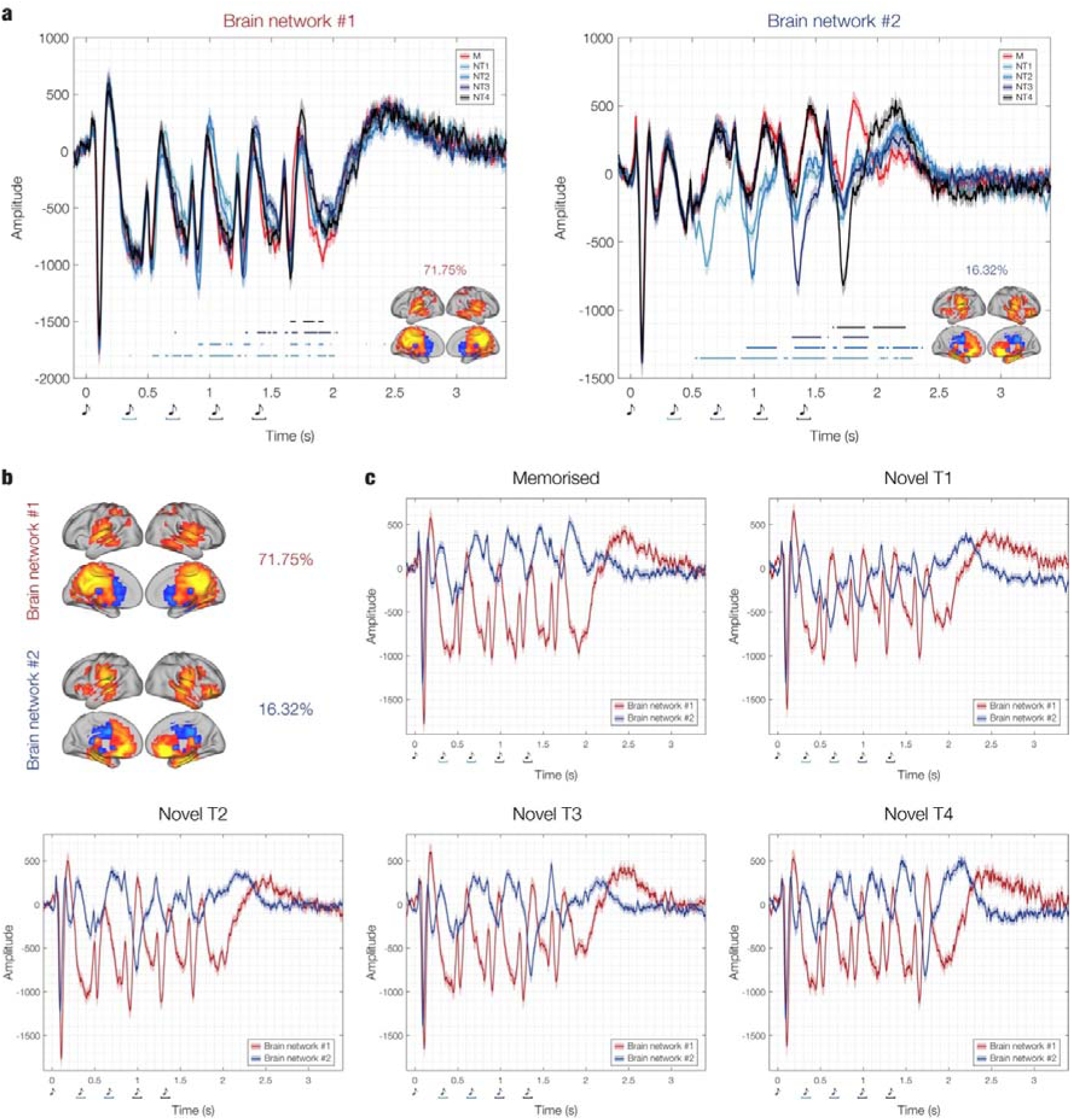
Time series and spatial activation patterns of the main brain networks. The figure illustrates the time series and spatial activation patterns of the two brain networks that explained the highest variance (71.75% and 16.32%, respectively). These networks were estimated using PCA within the BROAD-NESS framework, computed on the data averaged across conditions and participants. **a –** Independent time series for each participant, brain network, and experimental condition were generated using PCA-derived weights from the averaged data. The individual time series were then averaged across participants, as shown in the plots. Shaded areas represent standard errors. The brain templates illustrate the spatial extent of the networks, with yellow voxels contributing the most and light blue voxels contributing the least to the time series. Only voxels with values exceeding the mean by more than one standard deviation in absolute terms are depicted. Blue-black lines indicate the temporal extent of the significant differences between M versus each category of N (i.e. M versus NT1, M versus NT2, M versus NT3, M versus NT4), computed using two-sided t-tests corrected for multiple comparisons via False Discovery Rate (FDR). Different shades of blue-black represent specific M versus N comparisons. **b -** Focus on the spatial activation patterns of the brain networks. **c –** Focus on the brain networks time series, depicted independently for each experimental condition to emphasise similarities and differences between the two brain networks time series. Musical tone sketches represent the onset of each tone in the sequences.

The weights were subsequently multiplied by the original data to compute the time series for each brain network. It is important to note that while the weights were computed using data averaged across participants and conditions, the computation of the brain network time series was performed independently for each participant and experimental condition. This was necessary to allow statistical comparisons of the brain networks time series across experimental conditions. Specifically, statistical analyses were conducted by contrasting the time series of the memorised versus varied musical sequences, consistent with our prior study focusing on specific brain ROIs. Here, a two-sided t-test was performed for each time point and each combination of memorised versus novel sequences (i.e., M versus NT1, M versus NT2, M versus NT3, M versus NT4). Multiple comparisons were corrected using False Discovery Rate (FDR). This analysis revealed several significant time points where the memorised sequences differed significantly from the novel ones (**Figure 2**).

Detailed statistical results are summarised in **Table S1**.

### Data randomisations and brain networks

In addition to the MCS approach, we aimed to systematically assess how the sensitivity of BROAD-NESS was influenced by the temporal and spatial dimensions of latent brain networks. To achieve this, we independently disrupted the spatial and temporal organisation of the data matrix (brain voxels x time points) by implementing two types of randomizations. The randomisation strategies were designed as follows: (i) space randomisation, where the voxel indices were shuffled row-wise; and (ii) time randomisation, where the time indices were shuffled independently for each brain voxel. As in the case of the Monte Carlo simulations, to evaluate the stability and robustness of the procedure, each randomisation strategy was executed 100 times.

For the first randomisation (space randomisation), we found that the variance explained by the principal components and their associated time series remained identical to those of the original data (**Figure S4**). However, their spatial activation patterns exhibited a completely disrupted structure, indicating that this randomisation did not affect the PCA computation but resulted in non-meaningful spatial extents of the brain networks. In contrast, the second randomisation (time randomisation, which was also used within the MCS approach detailed above) disrupted both the spatial and temporal domains, leading to a significantly reduced variance explained by the main components (**Figure S4**). In this case, the combination of disrupted spatial activation patterns and time series displaying no differences between experimental conditions made the estimated brain networks not interpretable. The same statistical testing described for the original data was performed for both randomisation strategies. The time series and spatial activation patterns obtained from the two randomisation strategies are illustrated in **Figure 3**, while a complete statistical report is provided in **Table S2**.

**Figure 3:**
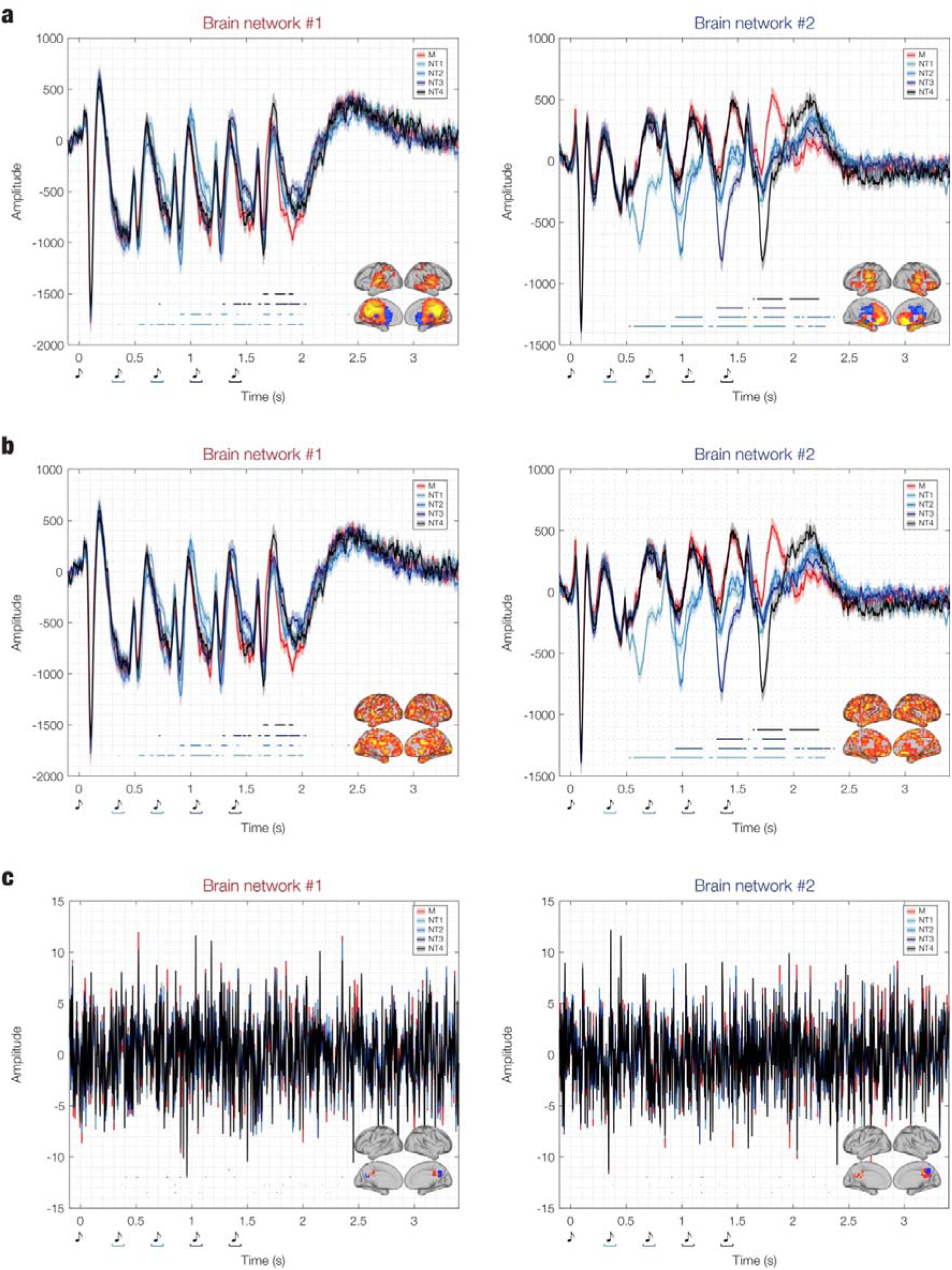
Brain networks estimated from randomised data. This figure illustrates the impact of data randomisation on the estimation of brain networks. **a -** Brain networks computed from the original data, identical to those shown in Figure 2. **b -** Space randomisation, where the order of the brain voxels was randomised without altering their time series. The time series and explained variance remain the same as in the original data, but the spatial activation patterns are scrambled. This demonstrates that PCA is independent of voxel order, confirming that meaningful patterns in the brain networks are genuine and not artefacts of the PCA algorithm. **c –** Time randomisation, where the time indices of the time series were disrupted, independently for each brain voxel. This leads to meaningless, low-amplitude time series and scrambled activation patterns, mostly concentrated around the mean. As a result, only a few brain voxels are depicted in the templates. In all cases, independent time series for each participant, brain network, and experimental condition were generated using PCA-derived weights from the averaged data. The individual time series were then averaged across participants, as shown in the plots. Shaded areas represent standard errors. The brain templates illustrate the spatial extent of the networks, with yellow voxels contributing the most and light blue voxels contributing the least to the time series. Only voxels with values exceeding the mean by more than one standard deviation in absolute terms are depicted. Blue-black lines indicate the temporal extent of the significant differences between M versus each category of N (i.e. M versus NT1, M versus NT2, M versus NT3, M versus NT4), computed using two-sided t-tests corrected for multiple comparisons via False Discovery Rate (FDR). Different shades of blue-black represent specific M versus N comparisons. The variance explained by the PCs for the original data and the two randomisation strategies is depicted in **Figure S4**.

### Comparing different PCA computations

To enhance the robustness of our analysis and elucidate the subtle differences arising from distinct PCA computation approaches to neural data, we conducted additional analyses. A key question relevant to experimental settings is whether it is more advantageous to compute PCA on individual experimental conditions or on aggregated data (e.g., data averaged over conditions).

In this study, we explored both approaches and compared the results, as illustrated in **Figure S5** and reported in detail in **Table S3**. Specifically, PCA was first computed on the data averaged across conditions, followed by the computation of time series. We also performed PCA independently for each condition, allowing for a direct comparison of the time series derived from the first PCs and an assessment of the differences. As shown in **Figure 4**, the two computations returned almost identical results, suggesting that the brain networks underlying the different experimental conditions were very similar (yet characterised by different temporal dynamics).

**Figure 4:**
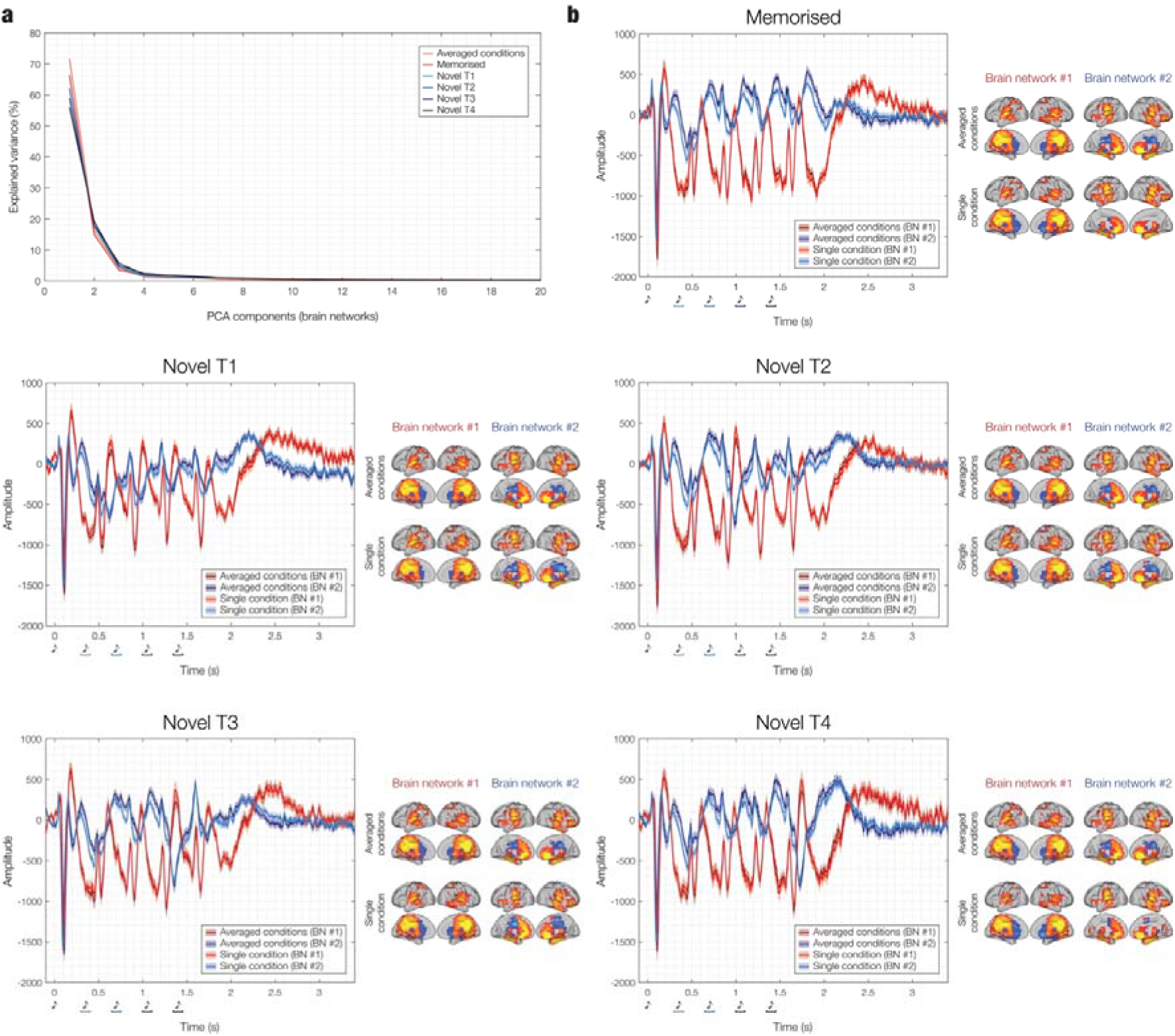
Brain networks estimated from data averaged across conditions or independently for each condition. **a -** Variance explained by the PCs after performing PCA on data averaged across conditions or independently for each condition. The nearly equivalent variance suggests that the different experimental conditions are linked to the same brain networks. **b -** Comparison of the time series for the two brain networks, computed using PCA weights from either averaged data or data selected independently for each condition. The time series are nearly identical, further supporting the notion that the brain networks underlying the different experimental conditions are extremely similar. Indeed, the differences between experimental conditions arise not in the brain networks spatial extent, but rather in the polarity and temporal dimension of their responses. In all cases, independent time series for each participant, brain network, and experimental condition were generated using PCA-derived weights from the data averaged across participants. The individual time series were then averaged across participants, as shown in the plots. Shaded areas represent standard errors. The brain templates illustrate the spatial extent of the networks, with yellow voxels contributing the most and light blue voxels contributing the least to the time series. Only voxels with values exceeding the mean by more than one standard deviation in absolute terms are depicted.

This concept can similarly be applied to the multiple participants involved in the study. PCA can be performed on the average data across participants, followed by the independent reconstruction of time series for each participant using the weights obtained from PCA. Alternatively, PCA can be computed independently for each participant, along with their respective time series, or it can be computed on the concatenated data from all participants and then the time series of each brain network can be reconstructed independently for each participant. We implemented all three approaches and presented a comparison in **Figure 5**, which revealed similar results, with the clearest and most interpretable outcomes obtained from PCA computed on the average data across participants.

**Figure 5:**
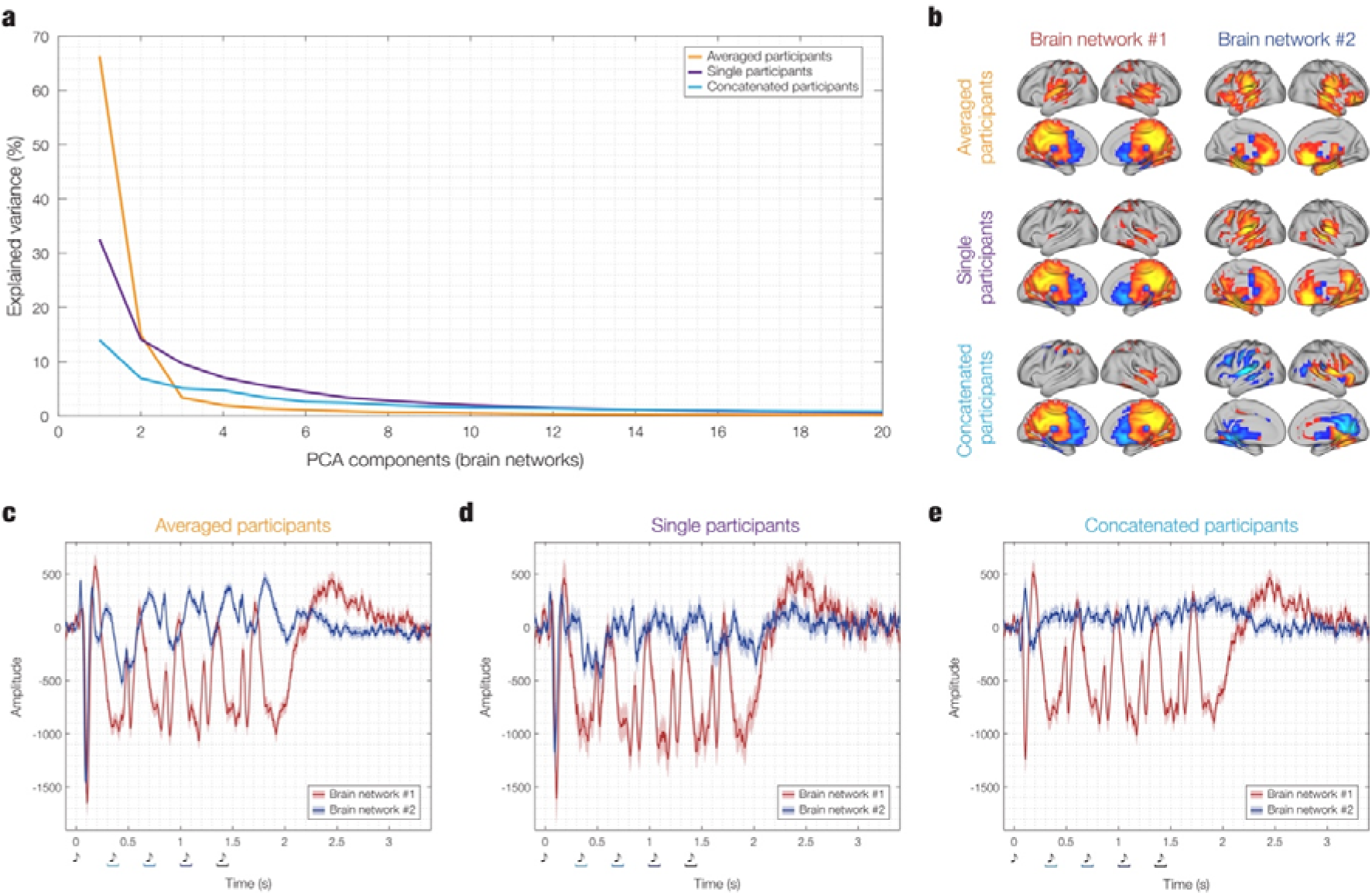
Brain networks estimated from data averaged across participants or independently for each participant. **a -** Variance explained by the PCs after performing PCA on data averaged across participants, or independently for each participant, considering one experimental condition: the previously memorised sequences. The results show a clear reduction in the variance explained by the first components when PCA is performed independently on individual participants (and the results averaged) or on the concatenated data from all participants, compared to PCA computed on data averaged across participants. **b -** The spatial activation patterns are depicted in brain templates, with yellow voxels contributing the most and light blue voxels the least to the corresponding time series. Only voxels with values exceeding the mean by more than one standard deviation in absolute terms are depicted. The plots show that the spatial activation patterns of the first brain network remain relatively stable, despite the drop in variance observed in **a**. However, the spatial activation patterns for the second brain network show a clear shift. When PCA is computed for individual participants (and results averaged), the network is less clearly defined but still maintains a similar spatial extent. In contrast, when PCA is performed on the concatenated data from all participants, the spatial extent of the second network drastically changes, losing its relevance. **c -** The time series were generated in three different ways. In the first case, independent time series for each participant and brain network were computed using PCA-derived weights from data averaged across participants. In the second case, PCA was computed independently for each participant, and individual time series were generated using participant-specific PCA weights. In the third case, PCA was computed on the concatenated data from all participants, and those weights were used to independently generate time series for each participant. In all cases, the individual time series were subsequently averaged across participants, as shown in the plots. Shaded areas represent standard errors. Overall, the time series results are consistent with the spatial activation patterns. The first brain network remains relatively stable across the different methods of computation, while the second network shows substantial changes. Specifically, the time series becomes less well-defined in the individual-participant computation and approaches a flat line in the concatenated data computation. This suggests that PCA should be performed on data averaged across participants, and the resulting weights should be used to compute the time series independently for each participant and experimental condition.

### Focus on multiple comparisons corrections

To further enhance the robustness of our results and provide additional insights into the contrasts between brain network time series while correcting for multiple comparisons, we computed and compared three additional correction methods specifically for the contrast between M and N1. The approaches, detailed in the methods section, include: (i) Bonferroni correction, (ii) cluster-based permutation test, and (iii) one-dimensional (1D) cluster-based Monte Carlo Simulation (MCS, α = .05, MCS *p-value* = .001). As illustrated in **Figure S5**, the results were highly convergent, confirming the effectiveness and consistency of all three correction methods.

## Discussion

Our study provides new insights into the large-scale brain networks underlying predictive coding during auditory memory recognition. Using the BROAD-NESS variant of the recently validated NESS framework^50^, we identified two distinct yet concurrent whole-brain networks involved in auditory and predictive processing, extending beyond conventional hierarchical models of forward and backward information flow. Crucially, while previous studies focused on predictive coding within limited brain regions or pairwise connections, we show that it operates at the level of whole-brain networks. These networks engaged auditory cortices in distinct interactions with other brain regions, exhibiting fine-grained spatial gradients and dynamic temporal patterns. Three key insights emerged from the current study.

First, we identified two primary whole-brain networks that were represented by the first two principal components, together explaining 88.06% of the variance (71.75% and 16.31%, respectively). Both networks prominently involved the auditory cortices but diverged in their broader connectivity and functional implications. The first network, also including the medial cingulate gyrus, captured early auditory processing, such as the N100 elicited by each sound, as well as later, slower ERF components sustaining activity between sounds. The second network, in addition to auditory cortices, involved hippocampal regions, the inferior temporal cortex, insula, and prefrontal regions, including the anterior cingulate and ventromedial prefrontal cortices. The time series of this second network indicated a slow positive ERF component for each sound, likely reflecting a process of matching incoming tones to stored memory traces of previous melodies^51,52^. Additionally, a more rapid negative peak emerged in response to variations in the sequence, which is consistent with prediction error responses^53,54^. These findings align with our previous ROI-based studies^32,33^, which identified hierarchical interactions between a restricted set of brain regions including auditory cortex, cingulate gyri, ventromedial prefrontal cortex, and hippocampus during the same auditory recognition task. The present findings also resonate with additional prior research. The network identified by the first principal component, involving the auditory cortices and cingulate gyrus, aligns with ROI-based studies on functional and structural connectivity, which have demonstrated strong links between these regions in both humans and animals^55–57^. This supports their joint role in early auditory processing and sustained attention^58–60^. Similarly, the network identified by the second principal component engaged regions typically associated with memory and prediction^61–63^. Previous studies have shown that these memory-related regions co-activate in both structural and functional connectivity, reinforcing their role in comparing incoming sounds with stored memory traces^64^. Crucially, unlike previous studies constrained to predefined ROIs, our findings reveal two whole-brain networks directly engaged in the task. Here, the auditory cortex appears to serve a dual function: engaging with medial cingulate structures for fundamental auditory processing while also interacting with memory-related regions for predictive computations. By leveraging the BROAD-NESS analytical framework, we reveal organisational principles of auditory memory recognition that were not accessible with ROI-based methods.

Second, our results relate to and extend the well-known dual-stream hypothesis, originally proposed for vision and later applied to auditory processing^65–68^. This framework posits two principal pathways: a ventral stream, leading from sensory areas to the medial temporal lobe and implicated in object recognition, and a dorsal stream, extending to the parietal lobe and primarily associated with spatial processing. Consistent with this hypothesis, one of the two networks identified in our study includes core regions of the ventral stream, such as the hippocampus, ventromedial prefrontal cortex, and inferior temporal cortex, supporting its role in recognition processes. Crucially, we demonstrate that this network not only aligns with the ventral stream but also exhibits clear evidence of predictive coding, encompassing both confirmed predictions and prediction errors unfolding over time. This suggests that predictive coding mechanisms are intrinsically linked with the role of the ventral stream in memory-based auditory processing. Interestingly, our findings also indicate a second, distinct network that differs from the classical dorsal stream. Instead of engaging the parietal lobe, this alternative network is centred around the auditory cortices and medial cingulate gyrus. While this network also exhibits predictive coding features, showing differences between experimental conditions and signal peaks in response to deviations in melodies, the predictive coding effects are notably weaker than in the ventral-stream-aligned network. Instead, this network appears to be more closely related to the general processing of auditory stimuli, including an early N100 ERF component followed by a slower negative peak, which shows only limited variation across conditions. These results suggest a novel perspective: rather than a direct dorsal counterpart, we identify a ’medial’ or ’central’ stream that supports sustained auditory processing and possibly attention, while playing a comparatively reduced role in the computation of confirmed prediction and prediction error.

Third, a major contribution of this study is the demonstration that predictive coding during auditory memory recognition is embedded within large-scale, orthogonal yet dynamically coordinated networks rather than being confined to local circuits or pairwise interactions (as typically observed by previous research in small networks formed by two-six ROIs). While previous research has primarily described predictive coding in terms of error propagation within restricted pathways^20,21^, our findings suggest that this process is integrated into broader network-level computations that simultaneously encode both confirmed predictions and deviations from expectation. This perspective shifts the focus from the conventional view of predictive coding as an exchange between discrete hierarchical levels toward a model in which distributed regions contribute in parallel to different aspects of prediction. The concurrent engagement of these networks implies that prediction and prediction error are not processed in isolation but unfold as part of a coordinated large-scale neural mechanism. Moreover, the differential involvement of these networks across time suggests that predictive coding operates across multiple temporal scales, integrating rapid sensory updates with slower memory-based processes. By revealing these organisational principles, our findings refine current theories of predictive processing in auditory system and memory^69,70^, suggesting that the brain’s predictive computations emerge from the occurrence and potential interplay of whole-brain networks rather than isolated functional modules.

While our study focused on the two primary networks that explained the majority of the variance, the BROAD-NESS analytical pipeline also revealed additional networks that were above chance level. Specifically, four smaller networks emerged, each accounting for a minor percentage of variance (4.16%, 1.30%, 0.99%, and 0.92%, respectively). These networks contained subsets of regions from the two primary networks but exhibited less interpretable activation patterns and time series. The less distinct temporal dynamics and activation patterns of these additional networks suggest that they likely reflect a minor proportion of residual activity mixed with noise, rather than major contributions to the predictive and memory processes.

Importantly, our findings on predictive coding for auditory sequences were enabled by the novel BROAD-NESS analytical pipeline, which represents a significant advancement over traditional methods in cognitive neuroscience for studying brain networks. Therefore, it is essential to discuss them also in relation to specific methodological aspects and nuances (for further technical details on BROAD-NESS, please refer to the Methods section and Supplementary Information). An essential remark is that traditional approaches to studying brain networks have typically relied on predefined ROIs, falling into three main categories: (i) pairwise connectivity without directionality (e.g. static functional connectivity^71^, instantaneous phase synchronisation^72^); (ii) pairwise connectivity with directionality (e.g. Granger causality^73^, transfer entropy^74^, partial directed coherence^75^); and (iii) functional network analyses constrained to predefined ROIs or parcellations (e.g. DCM^76^, graph theory on connectivity matrices^77^, hidden Markov models^78^). While these methods have yielded valuable insights, they are limited by their reliance on predefined brain regions, potentially overlooking key network dynamics. In contrast, BROAD-NESS operates on the entire source-reconstructed MEG space, analysing a high-resolution voxel grid (8 mm) that preserves spatial information while leveraging the high temporal resolution inherent to MEG. By avoiding ROI-based constraints^79–81^, this approach fully exploits the advantages of MEG source reconstruction, allowing for a data-driven exploration of the whole-brain networks. Furthermore, BROAD-NESS features a streamlined analytical pipeline that is both computationally efficient and based on only a few well-supported assumptions: (i) brain voxels are modelled as a linear combination of sources mixed with noise, with networks emerging when voxels covary over time; (ii) principal components derived from BROAD- NESS correspond to brain networks with interpretable dynamics; and (iii) variance explained by each component reflects the network’s relative significance. Moreover, BROAD-NESS is optimised for event-related designs, effectively isolating transient brain networks involved in stimulus processing. Altogether, these features enhance the interpretability of its outcomes, providing clear and straightforward insights into the brain networks underlying predictive coding in the auditory domain.

Beyond its immediate contributions, BROAD-NESS also provides a versatile framework for future research. For instance, although our approach does not explicitly assess non- linearity, BROAD-NESS can be integrated with methods such as transfer entropy or DCM to explore non-linear interactions at the network level. Additionally, although BROAD-NESS does not assume recurrent states, multivariate recurrence analysis can be applied to its time series to infer potential trajectories within a state-space defined by the orthogonal brain networks identified by BROAD-NESS.^82,83^. Similarly, directionality analyses can be performed on the identified network time series, offering further insights into causal interactions. These future directions highlight how BROAD-NESS not only provides novel insights in the current state, but also can be integrated with existing methodologies, paving the way for an even enriched understanding of the network dynamics underlying predictive coding.

In conclusion, our study provides novel insights into the large-scale network organisation of predictive coding in auditory memory recognition, revealing two concurrent whole-brain networks with distinct functional responses and roles. By leveraging the BROAD-NESS framework, we overcome previous methodological constraints and offer a fine-grained, interpretable characterisation of predictive processing at the whole-brain network level.

These findings not only refine our understanding of predictive coding but also open avenues for future research on its causal and dynamic properties across the whole brain.

## Methods

### Participants

The participant sample comprised 83 volunteers [33 males and 50 females (biological sex, self-reported)] aged between 19 and 63 years (mean age: 28.76 ± 8.06 years). Participants were recruited in Denmark, primarily from Western countries. Information on gender was not collected, as it fell outside the scope of this research. All participants were healthy, reported normal hearing, and had relatively homogeneous educational backgrounds. Specifically, 73 participants either held a university degree (bachelor’s or master’s, n = 54) or were currently enrolled as university students (n = 19). Of the remaining 10 participants, five held a professional degree earned after high school, and the other five had only completed high school. The study was approved by the Institutional Review Board (IRB) of Aarhus University (case number: DNC-IRB-2020-006) and conducted in accordance with the Declaration of Helsinki – Ethical Principles for Medical Research. All participants provided informed consent prior to the experiment and received compensation for their participation.

### Experimental stimuli and design

In this study, we employed an old/new auditory recognition task during magnetoencephalography (MEG) recordings (**Figure 1**), as described in previous studies**Error! Bookmark not defined.**,^103–107^. Participants first listened to a short musical piece twice and were asked to memorise it. The piece consisted of the first four bars of the right-hand part of Johann Sebastian Bach’s Prelude No. 2 in C Minor, BWV 847, where each bar contained 16 tones, totalling 64 tones. Each tone lasted approximately 350 ms, making the total duration 22,400 ms. A final tone, lasting 1000 ms, was added for a sense of closure, bringing the full duration to 23,400 ms (23.4 seconds). The musical notation for the prelude is shown in **Figure S1**.

After memorising the piece, participants were presented with 135 five-tone musical excerpts, each lasting 1750 ms. They were asked to determine whether each excerpt belonged to the original music (’memorised’ sequence [M], old) or was a variation (’novel’ sequence [N], new). Among the excerpts, 27 were taken directly from the original musical piece, while the remaining 108 were variations. **Figure S2** provides a visual representation of all the sequences used in the study.

The memorised sequences (M) were composed of the first five tones from the first three measures of the original piece, and each sequence was repeated nine times, totalling 27 trials. Novel sequences (N) were created by systematically varying the M sequences. For each of the three M sequences, nine variations were created by altering tones after the first (NT1), second (NT2), third (NT3), or fourth (NT4) tone, resulting in 27 variations per category and 108 novel sequences overall.

The variations followed several strategies. In some cases, the melodic contour was inverted, reversing the melodic direction of the original sequence. In other instances, the remaining tones were scrambled, or the same tone was repeated multiple times, sometimes varying only the octave. Another approach scrambled the intervals between tones. In most cases, the harmonic structure of the novel sequences was preserved relative to the original sequence. Further information about the sequences is reported in Bonetti et al.^33^.

This procedure allowed us to explore the brain networks dynamics involved in recognising previously memorised auditory sequences and in the conscious detection of sequence variations. The musical piece and sequences described above were generated as MIDI versions using Finale (version 26, MakeMusic, Boulder, CO) and presented through Psychopy v3.0.

### Data acquisition

The MEG recordings were conducted in a magnetically shielded room at Aarhus University Hospital (AUH), Denmark, using an Elekta Neuromag TRIUX MEG scanner with 306 channels (Elekta Neuromag, Helsinki, Finland). Data was collected at a sampling rate of 1,000 Hz with an analogue filtering of 0.1 - 330 Hz. Prior to the recordings, participants’ head shapes and the positions of four Head Position Indicator (HPI) coils were registered relative to three anatomical landmarks using a 3D digitizer (Polhemus Fastrak, Colchester, VT, USA). This information was later used to co-register the MEG data with anatomical MRI scans. Throughout the MEG recording, the HPI coils continuously tracked the position of the head, allowing for movement correction during analysis. Additionally, two sets of bipolar electrodes were used to monitor cardiac rhythm and eye movements, facilitating the removal of electrocardiography (ECG) and electrooculography (EOG) artefacts in the analysis phase.

MRI scans were obtained on a CE-approved 3T Siemens MRI scanner at AUH. The data included structural T1 (mprage with fat saturation) scans, with a spatial resolution of 1.0 x 1.0 x 1.0 mm and the following sequence parameters: echo time (TE) of 2.61 ms, repetition time (TR) of 2,300 ms, a reconstructed matrix size of 256 x 256, an echo spacing of 7.6 ms, and a bandwidth of 290 Hz/Px.

The MEG and MRI recordings were conducted on separate days.

### MEG data pre-processing

The raw MEG sensor data (comprising 204 planar gradiometers and 102 magnetometers) was initially pre-processed using MaxFilter^84^ (version 2.2.15) to reduce external interferences. We applied signal space separation (SSS) with the following MaxFilter parameters: downsampling from 1,000 Hz to 250 Hz, movement compensation via continuous HPI coils (default step size: 10 ms), and a correlation limit of 0.98 to reject overlapping signals between inner and outer subspaces during spatiotemporal SSS.

The MEG data was converted into Statistical Parametric Mapping (SPM) format and further processed and analysed using MATLAB (MathWorks, Natick, MA, USA), incorporating custom-built scripts (LBPD, https://github.com/leonardob92/LBPD-1.0.git) and the Oxford Centre for Human Brain Activity (OHBA) Software Library (OSL)^85^ (https://ohba-analysis.github.io/osl-docs/), which integrates Fieldtrip^86^, FSL^87^, and SPM^88^ toolboxes.

Next, the continuous MEG data underwent visual inspection via the OSLview tool to remove any large artifacts, though less than 0.1% of the collected data was discarded. Independent Component Analysis (ICA) was then used (with OSL implementation) to eliminate eyeblink and heartbeat artifacts^89^. The ICA process involved decomposes the original signal into independent components and correlating them with the activity recorded by the electrooculography (EOG) and electrocardiography (ECG) channels^90^. Components that showed a correlation at least three times higher than others were flagged as reflecting EOG or ECG activity. These flagged components were further validated through visual inspection, ensuring their topographic distribution matched typical eyeblink or heartbeat activity, and were subsequently discarded. The signal was then reconstructed using the remaining components.

Finally, the data was segmented into 135 trials (27 M, 27 NT1, 27 NT2, 27 NT3, and 27 NT4) and baseline-corrected by subtracting the mean baseline signal from the post-stimulus data. Each trial lasted 4500 ms (4400 ms of data and 100 ms of baseline).

### Source reconstruction

MEG is a powerful tool for detecting whole-brain activity with excellent temporal resolution. However, to fully understand the brain activity involved in complex cognitive tasks, it is essential to identify the spatial sources of this activity. This requires solving the inverse problem, as the MEG recordings reflect neural signals from outside the head but do not directly indicate which brain regions generated them. To address this, we employed beamforming algorithms^91,93,95^, using a combination of in-house-developed codes alongside the OSL, SPM, and FieldTrip toolboxes. The procedure consisted of two main steps: (i) designing a forward model and (ii) computing the inverse solution.

First, a single-shell forward model was constructed using an 8-mm grid. This head model treats each brain source as an active dipole and describes how the activity of such a dipole would be detected across MEG sensors^91,92^. The MEG data was co-registered with individual MRI T1-weighted images, using the 3D digitizer information to align the data. When individual anatomical scans were unavailable, an MNI152-T1 template with 8-mm spatial resolution was used to compute the forward model.

Second, a beamforming algorithm was applied as the inverse model. Beamforming uses a set of weights applied to different source locations (dipoles) to isolate the contribution of each brain source to the activity recorded by the MEG sensors^93–95^. This was done for every time point of the recorded brain data, allowing us to reconstruct the spatial sources of the MEG signal. The detailed steps for implementing the beamforming algorithm are provided below. The data recorded by the MEG sensors (*B*) at time *t*, can be described by the following Eq. (1):

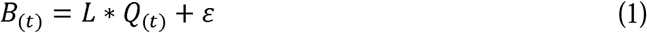

where *L* is the leadfield model, *Q* is the dipole matrix carrying the activity of each active dipole (*q*) at time *t*, and LJ is noise (for details, see Huang et al.^91^) To solve the inverse problem, *Q* must be estimated for each *q*. In the beamforming algorithm, a series of weights are computed to describe the transition from MEG sensors to the active dipole *q*, independently for each time point. This is reported in Eq. (2):

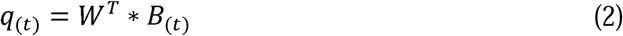

Here, the superscript *T* refers to transpose matrix. To compute the weights (*W*), matrix multiplication between *L* and the covariance matrix of MEG sensors (*C*) is performed. Importantly, the covariance matrix *C* was computed on the signal after concatenating the single trials of all experimental conditions. For each dipole *q*, the weights (*W_q_*) were computed as shown in Eq. (3):

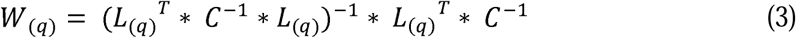

The computation of the leadfield model *L* was performed for the three main orientations of each dipole ^92^. Before computing the weights, to simplify the beamforming output^96,97^, the orientations were reduced to one using the singular value decomposition algorithm on the matrix multiplication reported in Eq. (4):

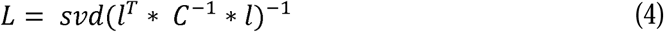

Here, *l* represents the leadfield model with the three orientations, while *L* is the resolved one-orientation model that was utilized in Eq. (3).

Once computed, the weights were normalised for counterbalancing the reconstruction bias towards the center of the head ^93,98–100^ and applied to the neural activity averaged over trials, independently for each time point (Eq. 2) and experimental condition. This procedure returned a time series for each of the 3,559 brain sources^93,99^.

### Principal Component Analysis (PCA), Monte-Carlo simulations (MCS) and randomisations

PCA was applied to the 3,559 reconstructed brain sources time series to disentangle broadband brain networks operating simultaneously. PCA is a dimensionality reduction method that transforms data into new variables, or principal components, which account for the most variance within the dataset. This is achieved by computing the eigenvectors and eigenvalues of the data’s covariance matrix and then projecting the data onto the directions of maximum variance. Traditionally, PCA simplifies high-dimensional data while retaining the most important information.

In this study, however, PCA was applied to a dense set of MEG-reconstructed brain data at the voxel level (3,559 brain voxel time series). The goal was not just to reduce the dimensionality but to identify brain networks (corresponding to the PCs) through the application of PCA on the 3,559 brain voxel time series.

In short, PCA operates as follows: First, the data is centred by subtracting the mean of each brain voxel timeseries from the dataset *X*, as represented by Eq. (5):

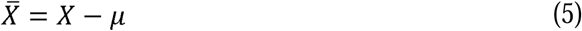

where µ is the mean vector of *X*.

Then, the covariance matrix *C* is computed on the centred data, as shown in Eq. (6):

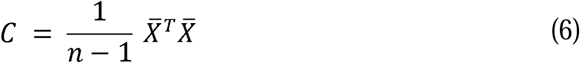

where *n* is the number of data points.

Then the eigenvalue equation for the covariance matrix is solved to find eigenvalues and eigenvectors, Eq. (7):

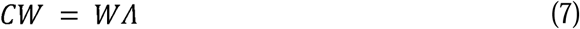

Where W is the eigenvector matrix (set of weights for each principal component) and A is the diagonal matrix of the corresponding eigenvalues (which indicate the amount of variance explained by each component).

Then, the eigenvectors w were used to compute the activation time series of each brain network *Y* by multiplying them by the original data *X*, as shown in Eq. (8):

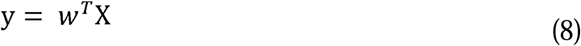

Furthermore, we computed the spatial projection of the components (spatial activation patterns *A*) in brain voxel space. This was done by multiplying the weights of the analysis (the eigenvectors w in this case) by the covariance matrix *C*, as shown in Eq. (9):

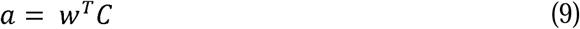

Interestingly, while previous studies on interpreting weights from multivariate analyses in MEG data have recommended calculating spatial activation patterns^101^, in this case, the relative contribution of each brain voxel to the network remains the same, whether using the direct eigenvector w or computing the spatial activation patterns.

Importantly, as detailed in the following section, the PCA procedure was applied across various scenarios to ensure a comprehensive evaluation of the algorithm’s performance. This included performing PCA independently for each participant and condition, as well as on data averaged across participants and conditions.

Finally, we aimed to systematically assess how the sensitivity of BROAD-NESS was influenced by the temporal and spatial dimensions of latent brain networks. To achieve this, we independently disrupted the spatial and temporal organisation of the data matrix (brain voxels x time points) by implementing two types of randomizations. The randomisation strategies were designed as follows: (i) space randomisation, where the voxel indices were shuffled row-wise; and (ii) time randomisation, where the time indices were shuffled independently for each brain voxel. To evaluate the stability and robustness of the procedure, each randomisation strategy was executed 100 times. The same statistical testing described for the original data was also performed for both randomisation strategies.

Additionally, the second randomisation was employed in a Monte Carlo simulation (MCS) solution. In this case, we retained the variance explained by the first component of the PCA computed on the randomised data for each of the 100 permutations. This produced a reference distribution of the variance explained by the first component in the randomised data. This reference distribution was then compared to the variance explained by the components in the original data, which were deemed significant only if they explained a greater variance than the highest explained variance in the reference distribution.

Time series and spatial activation patterns of the brain networks that were significant following the MCS approach are illustrated in **Figures 2** and **S3**, while the selective effects of each randomisation strategy are presented in **Figures 3** and **S4**.

### Comparing different PCA computations

To enhance the robustness of our work and provide greater insight into the subtle differences associated with the computation of PCA on neural data, we examined a crucial aspect relevant to experimental settings. Specifically, when multiple experimental conditions are present, it is essential to determine whether it is more beneficial to compute PCA on individual conditions or on aggregated conditions (e.g., data averaged over conditions).

To address this, we computed both approaches and compared the results, as illustrated in **Figure 4**. Specifically, we first conducted PCA on the data averaged across conditions and then reconstructed the time series. Subsequently, we computed PCA independently for each condition. This allowed us to compare the brain networks time series of the conditions computed in the two approaches and assess any difference.

A similar consideration applies to the multiple participants in our study, where we explored three possible approaches for computing PCA. The first approach involved performing PCA on the average across participants, followed by reconstructing the time series independently for each participant using the weights obtained from this analysis. The second approach entailed performing PCA independently for each participant along with their respective time series. Finally, the third approach consisted of computing PCA on the concatenated data from all participants, and then deriving the time series independently for each participant.

We implemented all three approaches and compared the results, as illustrated in **Figure 5**.

### Statistical analysis

After computing PCA and the time series of the brain networks for each participant and experimental condition, we conducted statistical analyses by contrasting the time series of the previously memorised versus the varied melodies. This approach was consistent with our earlier study that focused solely on brain ROIs^33^. Specifically, we performed a two-sided t-test for each time point across all combinations of M versus N melodies (i.e., M versus NT1, M versus NT2, M versus NT3, M versus NT4). Subsequently, we applied False Discovery Rate (FDR) corrections to account for multiple comparisons.

To strengthen the robustness of our findings and offer deeper insights into the process of contrasting time series of brain networks, we implemented and compared three alternative methods for correcting for multiple comparisons. This analysis specifically focused on the contrast between M versus NT1, as illustrated in **Figure S5**.

First, we applied the Bonferroni correction, which is a stringent method that adjusts the significance level (a = .05) by dividing it by the number of tests performed, thereby reducing the likelihood of false positives.

Second, we used a cluster-based permutation test, as described by Maris and Oostenveld^102^. Initially, we conducted statistical tests on the original data, as described earlier. We then identified clusters of neighbouring significant time points by applying a threshold to the statistical results (α = .05). Next, we performed 1000 permutations, where, for each permutation, the labels of the two experimental conditions were shuffled for each participant, and the statistics were recalculated. For each permutation, we computed the maximum cluster size of the significant time points, mirroring the approach used for the original data. This yielded a reference distribution of significant clusters, which we compared with the clusters from the original data. Clusters in the original data were deemed significant only if their sizes exceeded the maximum cluster size observed in the permuted data at least 99.9% of the time.

Third, we conducted a one-dimensional (1D) cluster-based MCS (MCS, α = .05, MCS p-value = .001), as detailed in our previous studies^32,33,103–109^. This approach involved identifying clusters of significant neighbouring time points obtained from the t-tests described above and then performing 1000 permutations to randomise these significant values. For each permutation, we extracted the maximum cluster size to establish a reference distribution. The original clusters were considered significant if they were larger than 99.9% of the clusters obtained from the permuted statistical results. This method is similar to the cluster-based permutation test but should be regarded as an empirical approach. Its primary advantage lies in its computational efficiency, delivering results much faster than the permutation test as it does not require recalculating statistics. Instead, it works directly with clusters formed from the statistics computed on the original data. However, a limitation is that this empirical method does not adhere to classical inferential statistical principles. In our analysis, we clarify how this approach relates to the permutation test, highlighting their respective advantages and limitations in the context of statistical testing for brain data time series.

## Data and code availability

The pre-processed neuroimaging data generated in this study have been deposited in the Zenodo database under accession code [https://doi.org/10.5281/zenodo.10715160]110 and are publicly available.

The MEG data was first pre-processed using MaxFilter 2.2.15. Then, the data was further pre-processed using Matlab R2016a or later (MathWorks, Natick, Massachusetts, United States of America). Specifically, we used codes from the Oxford Centre for Human Brain Activity Software Library (OSL), FMRIB Software Library (FSL) 6.0, SPM12 and Fieldtrip.

In-house-built code and functions used in this study are part of the LBPD repository which is available at the following link: https://github.com/leonardob92/LBPD-1.0.git

The full analysis pipeline used in this study is available at the following link: https://github.com/leonardob92/BROADNESS_MEG_AuditoryRecognition.git

## Acknowledgements

The Center for Music in the Brain (MIB) is funded by the Danish National Research Foundation (project number DNRF117).

L.B. is supported by Lundbeck Foundation (Talent Prize 2022), Carlsberg Foundation (CF20-0239), Center for Music in the Brain, Linacre College of the University of Oxford and Nordic Mensa Fund.

M.L.K. is supported by Center for Music in the Brain and Centre for Eudaimonia and Human Flourishing, which is funded by the Pettit and Carlsberg Foundations.

In addition, we thank the Department of Physics of the University of Bologna for the support provided to C.M., Fundación Mutua Madrileña (Mutua Madrileña Foundation, Madrid, Spain) for the support provided to G.F.R. and Mensa: The International High IQ Society (Italian section) for the support provided to F.C.

## Author contributions statement

L.B., G.F.R., M.H.A. and M.R. conceived the hypotheses. L.B. designed the study. L.B., M.L.K., C.T. and P.V. recruited the resources for performing data collection and analysis. L.B., G.F.R. and F.C. collected the data. L.B., G.F.R., M.H.A., C.M. and M.R. performed pre-processing, statistical analysis and linear decomposition of the neural signal. L.B., C.M., M.H.A. and M.R. developed the code for the BROAD-NESS framework. M.L.K., C.T., P.V. and M.R. provided essential help to interpret and frame the results within the neuroscientific and analytical literature. M.H.A. and L.B. wrote the first draft of the manuscript, which was primarily integrated by M.R. and G.F.R. The figures were prepared by L.B., G.F.R. and C.M. All the authors contributed to and approved the final version of the manuscript.

